# Alzheimer’s disease-associated genotypes differentially influence chronic evoked seizure outcomes and antiseizure medicine activity in aged mice

**DOI:** 10.1101/2024.10.06.616921

**Authors:** Kevin M. Knox, Stephanie Davidson, Leanne M. Lehmann, Erica Skinner, Alexandria Lo, Suman Jayadev, Melissa Barker-Haliski

## Abstract

**INTRODUCTION:** Alzheimer’s disease (AD) patients are at greater risk of focal seizures than similarly aged adults; these seizures, left untreated, may worsen functional decline. Older people with epilepsy generally respond well to antiseizure medications (ASMs). However, whether specific ASMs can differentially control seizures in AD is unknown. The corneal kindled mouse model of acquired chronic secondarily generalized focal seizures allows for precisely timed drug administration studies to quantify the efficacy and tolerability of ASMs in an AD-associated genetic model. We hypothesized that mechanistically distinct ASMs would exert differential anticonvulsant activity and tolerability in aged AD mice (8-15 months) to define whether rational ASM selection may benefit specific AD genotypes.

**METHODS:** Aged male and female PSEN2-N141I versus age-matched non-transgenic control (PSEN2 control) C57Bl/6J mice, and APP^swe^/PS1^dE9^ versus transgene negative (APP control) littermates underwent corneal kindling to quantify latency to fully kindled criterion. Dose-related ASM efficacy was then compared in each AD model versus matched control over 1-2 months using ASMs commonly prescribed in older adults with epilepsy: valproic acid, levetiracetam, lamotrigine, phenobarbital, and gabapentin.

**RESULTS:** Sex and AD genotype differentially impacted seizure susceptibility. Male PSEN2-N141I mice required more stimulations to attain kindling criterion (X^2^=5.521; p<0.05). Male APP/PS1 mice did not differ in kindling rate versus APP control mice, but they did have more severe seizures. There were significant ASM class-specific differences in acute seizure control and dose-related tolerability. APP/PS1 mice were more sensitive than APP controls to valproic acid, levetiracetam, and gabapentin. PSEN2-N141I mice were more sensitive than PSEN2 controls to valproic acid and lamotrigine.

**DISCUSSION:** AD genotypes may differentially impact ASMs activity and tolerability in vivo with advanced biological age. These findings highlight the heterogeneity of seizure risk in AD and suggest that precisely selected ASMs may beneficially control seizures in AD, thus reducing functional decline.

## Background

Focal impaired awareness seizures are increasingly recognized to co-occur in Alzheimer’s disease (AD) and other dementias. These untreated seizures may accelerate onset of neuropsychiatric symptoms and worsen cognitive decline^1; 2^. This is particularly tragic because antiseizure medicines (ASMs) are widely available, safe, and effective in mitigating seizure burden in people with epilepsy.

Despite the greater incidence of epilepsy in older adults and increased risk of comorbid seizures in AD, few studies have differentiated the potency and tolerability of FDA-approved ASMs in aged animal seizure models^3; 4^. Such information could inform potential risk for tolerability and drug-drug interactions in older adults with seizures, as well as advance precision medicine discovery platforms for disease-modifying therapeutics for AD^5^. ASMs on the market were largely brought forth based on efficacy in young adult, neurologically-intact rodents^6^ without considering genetic risk factors relevant to AD. Preclinical information concerning the efficacy and safety of promising investigational ASMs is infrequently evaluated in aged animals with chronic seizures^6; 7^. Few studies assess ASMs in rodents with AD-associated variants, thus our study objective was to inform precision medicine selection for seizures in AD.

Autosomal dominant early-onset AD is associated with variants in three genes: amyloid precursor protein (APP), the homologous presenilin 1 (*PSEN1)* and presenilin 2 (*PSEN2*) proteins^8; 9^. Presenilin (PS) proteins are intramembrane proteases of the catalytic component of γ-secretase, mutations of which can shift APP cleavage products towards the neurotoxic fragment of amyloid, Aβ^1–4210^. However, *PSEN* variants actually reduce overall proteolytic activity, thereby not directly increasing Aβ protein aggregates^11^. Nonetheless, these three risk genes confer nearly 100% penetrance, making models harboring these genetic variants relevant to study AD and its comorbidities, including seizures^12^. AD patients with the most common *PSEN2* variant (N141I) exhibit a high incidence of comorbid seizures^13^. Within 5 years of AD diagnosis, over 28% of patients with *PSEN2* mutations, and 31% of patients with APP duplications reported at least one seizure^9^. Aged mice expressing these AD-associated risk factors may be thus applied to quantify ASM activity and tolerability to inform precise seizure management strategies in clinical AD.

There is differential susceptibility to chronic neuronal network hyperexcitability in young adult APP-overexpressing^12; 14^ and *PSEN2* transgenic^14^ or knockout(KO)^15; 16^ mice suggesting that AD genotypes do not uniformly influence seizure susceptibility. Loss of normal *PSEN2* function is associated with dramatic, age-related shifts in susceptibility to epileptogenesis and ASM efficacy^15; 16^. However, it is currently unknown whether biological age and AD-associated genotypes change seizure risk or ASM activity. Use of well-characterized seizure and epilepsy models in AD-associated transgenic mice thus offers the potential to uncover novel molecular contributors underlying hyperexcitability and epileptogenesis across a lifespan, which may potentially reveal targeted anticonvulsant drugs specifically for this patient group. This present study was thus designed to assess the chronic seizure susceptibility in two aged AD-associated models, PSEN2-N141I mice^17^ expressing only the human *PSEN2* variant, and APP^Swe^/PSEN1^dE918^ double mutant mice that develop severe amyloidopathy. Various AD-associated mouse models demonstrate differential susceptibility to seizures in early life (2-4 months of age)^12; 14; 15; 19; 20^, but few studies have conducted such assessments in aged rodents^15^. We then sought to validate a drug discovery paradigm to determine whether mechanistically distinct ASMs differentially influence seizure control in an AD-associated aged brain. The secondary objective of this project was to therefore assess the Aβ-dependent and independent contributions to ASM activity, uncovering precision treatment strategies for the comorbid seizures of AD. We herein show novel age-related Aβ-dependent (APP/PS1) and Aβ-independent mechanisms (PSEN1-N141I) underlying seizure risk and ASM activity. This study informs clinical trial design and rational ASM selection in AD patients with epileptiform activity^21^.

## Methods

### Animals

Mice were housed on a 14:10 light cycle (on at 6 h00; off at 20 h00) in ventilated cages with free access to food and water,^22^ and as approved by the UW Institutional Animal Care and Use Committee (protocol 4387-01), conforming to ARRIVE guidelines^23^. All testing was performed blinded to genotype.

### Antiseizure Medicines

ASMs were administered by the intraperitoneal (i.p.; Table 1) route and tested consistent with our ASM screening approach in wild-type (WT) mice^7; 24^. All ASMs were formulated in 0.5% methylcellulose (Sigma Aldrich #M0430, St. Louis, MO USA).

**Table 1.**
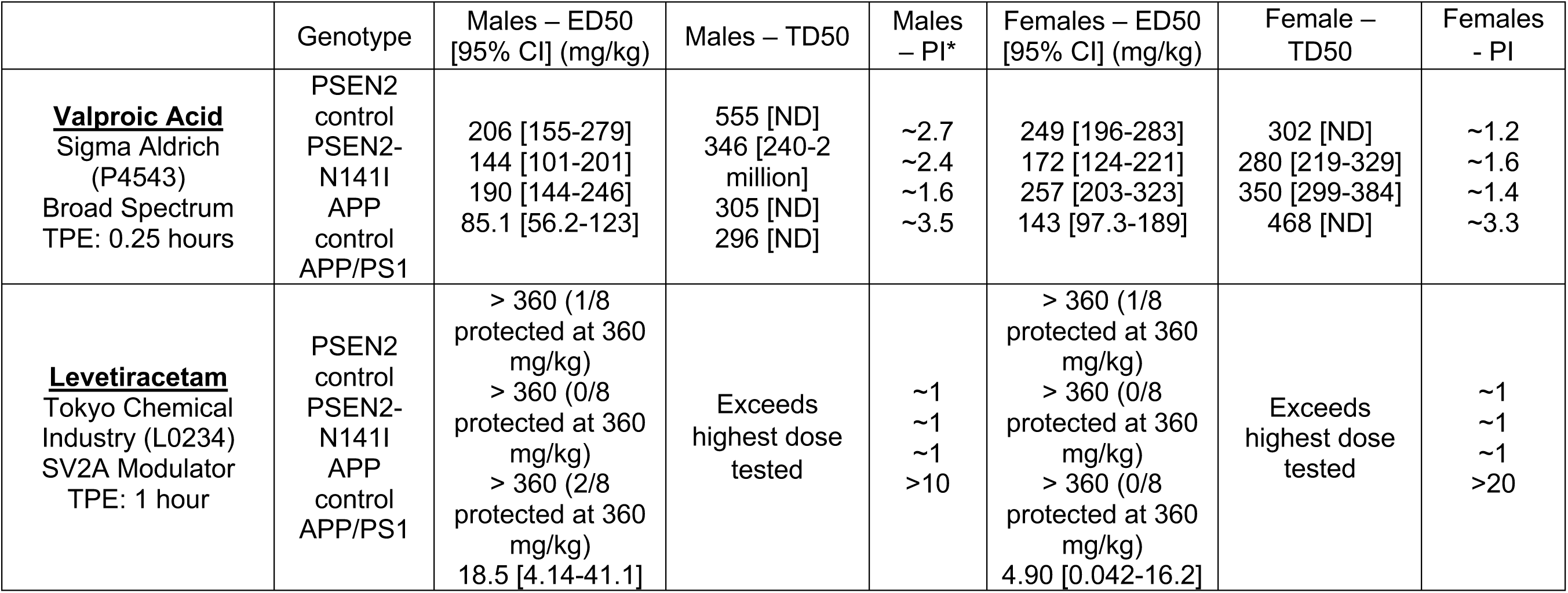
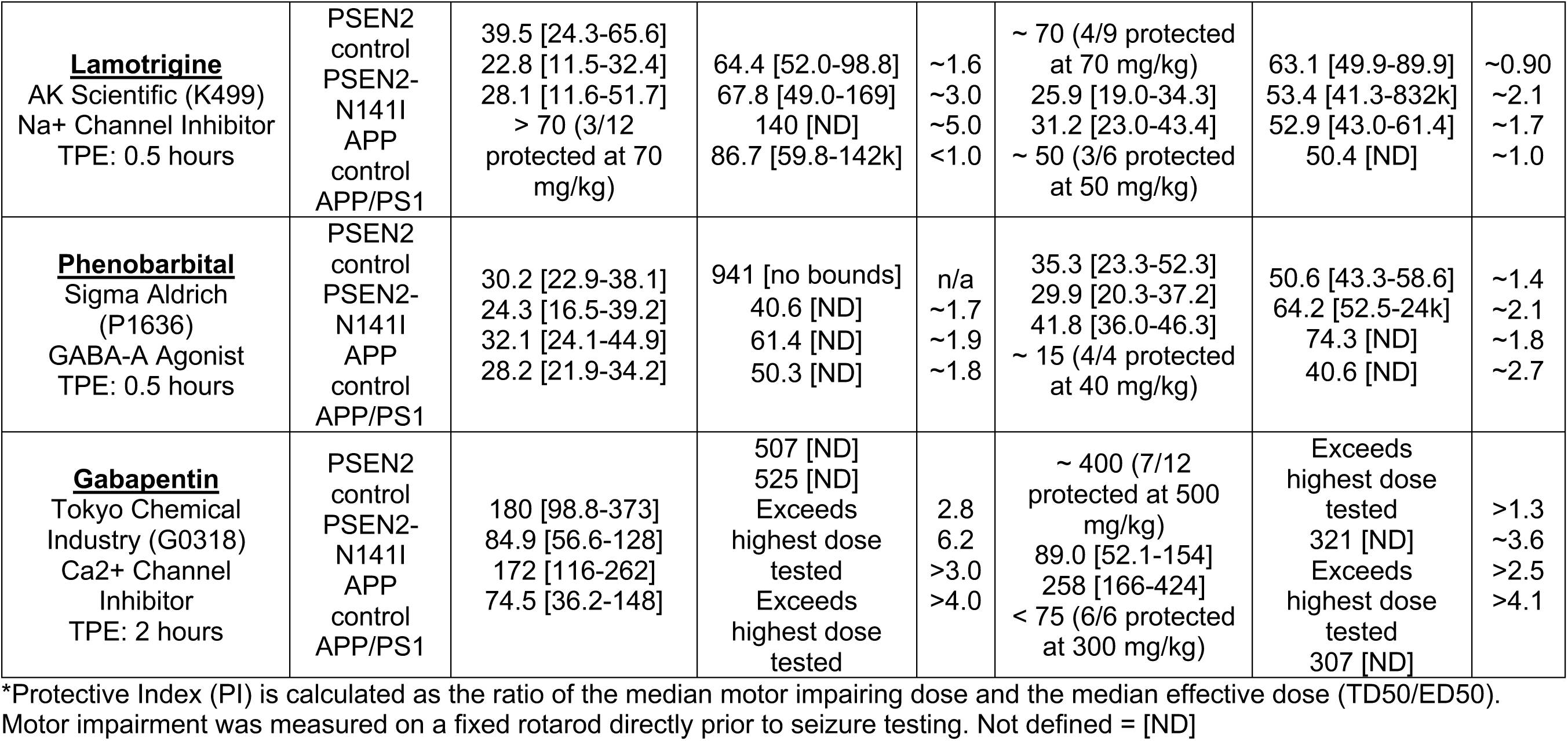
Calculated median effective (ED50) and motor impairing dose (TD50) in male and female aged transgenic control and AD mouse models.

### Corneal Kindling

60 Hz corneal kindling was evoked with an initially benign electrical current (3 sec, 60 Hz, sinusoidal pulse; 3.0 mA for males or 2.6 mA for females)^14; 15; 18; 24; 25^. Any mouse not achieving kindling criterion was excluded from ASM profiling.

Six cohorts of aged mice (8-15 months-old at initiation) established the dose-related ASM response (Figures 3 and 4). Male mice group sizes for pharmacology testing are detailed in Supplemental Table 1. An additional aged mice cohort was used for neuropathological assessment following kindling or sham kindling (Figure 5).

### Seizure Duration

Behavioral kindled seizure duration was recorded from all fully kindled mice following a 3-5-day stimulation-free period after kindling acquisition, as previously described^15^.

### Antiseizure Medication Dose-Response Studies in Fully Kindled Mice

Repeated ASM testing commenced at least 5-7 days after achieving kindling criterion. Seizure scores ≥ 2 were “protected”. Mice were administered escalating doses of each ASM bracketing previously established median effective doses (ED50) in kindled male WT mice^24^. Mice were allowed a minimum of 3 days’ washout between each ASM administration in a cross-over design to account for drug and seizure history^15; 24; 25^.

### Cryosectioning for Immunofluorescence

*–* Mice for immunofluorescence were euthanized 24 hours after achieving kindled criterion, with brains frozen and cryosectioed (Leica DM1860), as previously described^20; 26^.

### Amyloid beta and astroglia immunofluorescence

– One slide/mouse was processed for Aβ (6e10) and astrocytes (GFAP) labeling as previously reported^14^. An additional slide was processed for the presence of Aβ plaques (Thioflavin S). Photomicrographs were captured with a fluorescent microscope (Leica DM-4) with a 20x objective (80x final magnification), with acquisition settings held constant. We quantified the presence of positive labeling with both 6e10 and Thioflavin S in dorsal hippocampal structures (CA1, CA3, and dentate gyrus), and cortex directly overlying dorsal hippocampus (yes/no with positive signal) by two blinded investigators (Table 2).

**Table 2.**
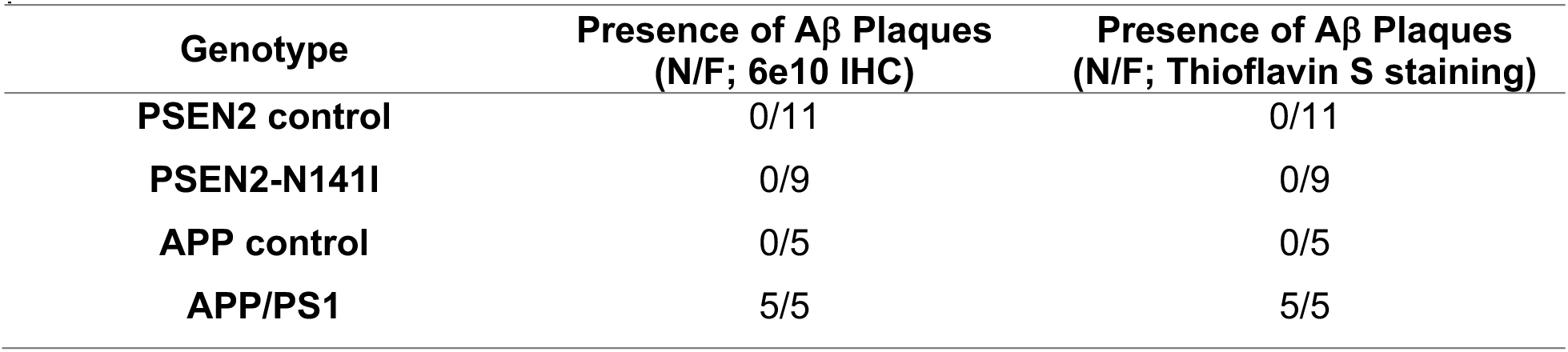
Qualitative assessment of dense core Aβ plaques in sham and fully kindled male and female APP/PS1 and PSEN2-N141I versus control mice indicates that only aged APP/PS1 mice demonstrated plaq<u>ue accumulation.</u>

### Statistics

– Latency to kindling criterion (Figure 1) and premature mortality (Figure 2) were quantified by a log-rank test and presented as Kaplan Meier survival plots. Seizure duration was quantified by a one-way ANOVA (genotype) within sex (Figure 2). Body weight changes during the chronic ASM administration period were quantified by two-factor ANOVA. All statistical analysis was conducted in GraphPad Prism v9.0 or later, with p<0.05 significant. When possible, an ED50 and median impairing dose (TD50) for each ASM was calculated by Probit regression (XLStat; Addinsoft)^27^, with protective index (PI) calculated (ratio of TD50:ED50; Table 1).

**Figure 1.**
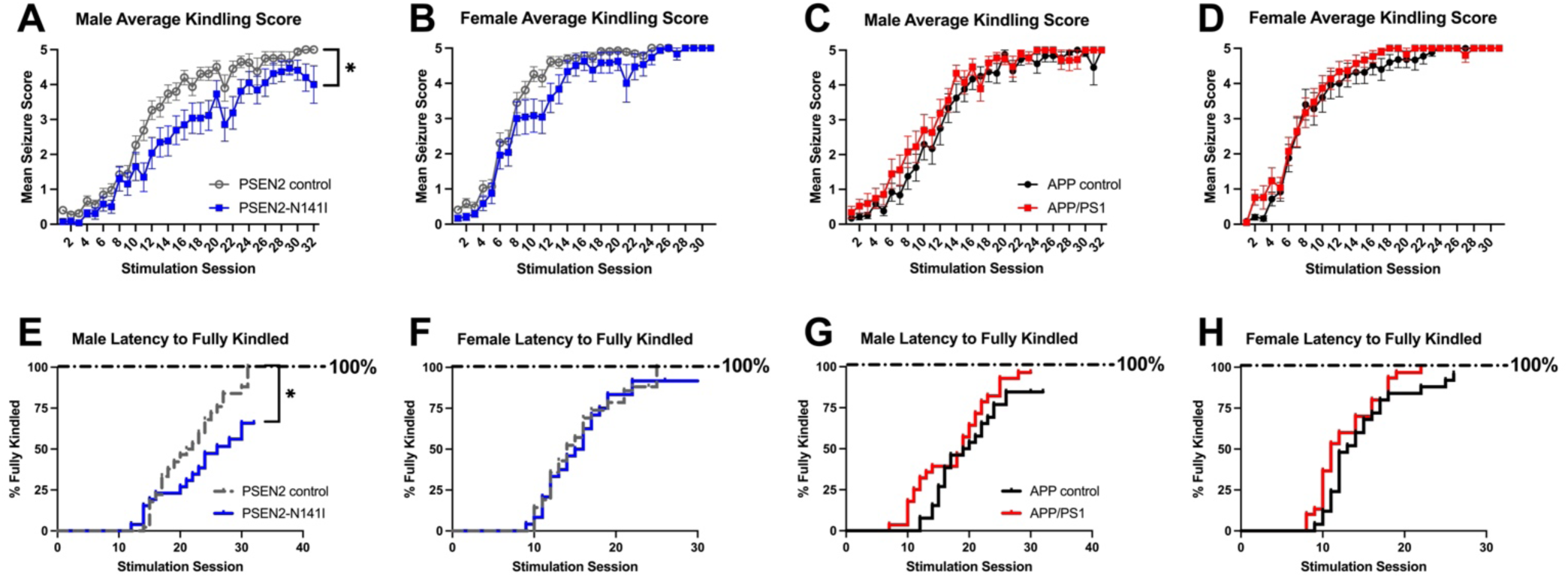
Corneal kindling acquisition curves for aged (8-15 months old) PSEN2 control and PSEN2-N141I (A) male and (B) female mice; APP control and APP/PS1 (C) male and (D) female mice. Data for graphs A-D are presented as the average Racine stage seizure score for each genotype at each separate stimulation session. A) Aged male PSEN2-N141I mice kindled significantly slower than aged-matched control mice. C) Further aged APP/PS1 male mice did not kindle at any different rate versus age-matched control mice. Aged female B) PSEN2-N141I and D) APP/PS1 mice do not acquire the fully kindled state at any different rate relative to age-matched control females. The latency to attain the fully kindled state (E-H) was measured as the number of stimulation sessions required for a mouse to have 5 consecutive stage 5 seizures. Aged PSEN2-N141I transgenic male mice (E) reached kindling criterion slower than PSEN2 control mice. There was a difference in the latency to the fully kindled state in male APP/PS1 mice (G) compared to APP control. There were no marked differences latency to attain kindling criterion in aged female PSEN2-N141I (F) or APP/PS1 (H) mice.

**Figure 2.**
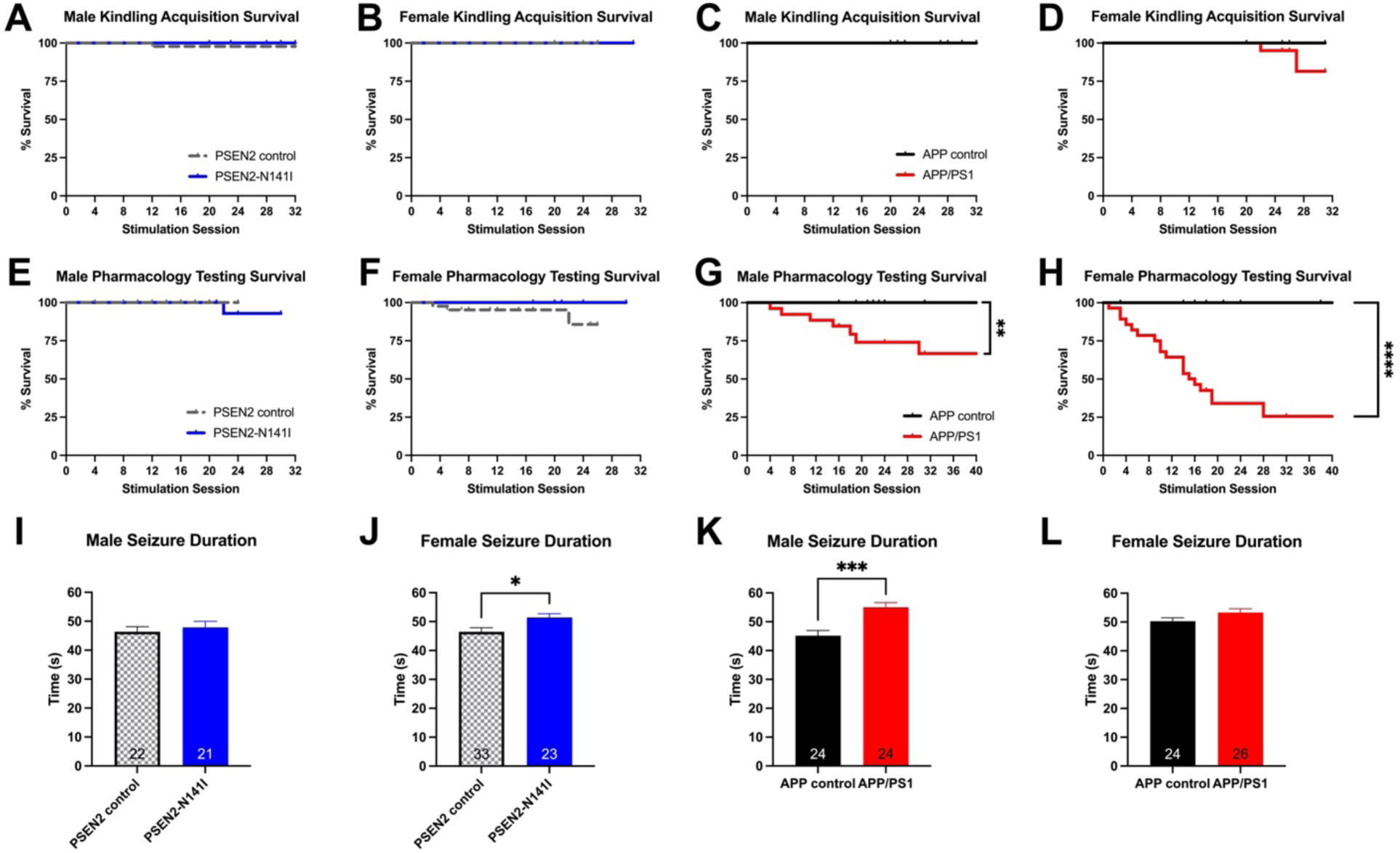
Induction of the 60 Hz corneal kindling model in aged male and female mice with AD-related genotypes leads to differences in seizure-induced survival outcomes. (A-D) There was no significant difference in the survival rate of animals of any genotype or sex during the 3 week, twice-daily electrical kindling period. Once animals attained the fully kindled state, they were enrolled for subsequent ASM screening twice/week for 1-2 months, or until spontaneous death. There was no effect of genotype on mortality during repeated ASM testing in either PSEN2-N141I male (E) or female (F) mice. Notably, both male (G) and female (H) APP/PS1 mice with kindled seizures experienced significant mortality during the ASM testing period. Between kindling acquisition and pharmacology testing upon attaining the fully kindled state, the duration of a single evoked seizure was assessed in all animals following a 5-7 day stimulation-free period. PSEN2-N141I male mice (I) showed no change in seizure duration from control mice while PSEN2-N141I female mice (J) had a significantly longer seizure duration than their controls. Notably, male APP/PS1 mice (K) had significantly longer seizures than their controls while the females (L) showed only a trend towards longer seizures.

## Results

### AD-related genotypes differentially influence kindling in aged mice in a sex-related manner

We have established that young adult mice with AD-associated genotypes exhibit stark differences in kindling acquisition rate with APP/PS1 mice having accelerated kindling and PSEN2 transgenic models having a delay in kindling^14; 15; 18^. Our present study extends these earlier findings to demonstrate that aged (8-15 month-old) male PSEN2-N141I mice require significantly more stimulations to reach criterion (X^2^=5.03; p=0.25) versus age-matched PSEN2 controls (Figure 1A, 1E). There was no change in reaching kindling criterion for female PSEN2-N141I mice (Figure 1B, 1F). Kindling rate was no different between male (Figure 1C, 1G) or female (Figure 1D, 1H) APP/PS1 mice and APP controls. Therefore, both age and AD-associated genotype significantly influenced kindling acquisition and hyperexcitable neuronal network formation.

### Evoked chronic focal seizures accelerate mortality in aged APP/PS1 mice

Chronic kindled seizures are generally well-tolerated in male and female WT mice^28^. Yet we have earlier reported that kindled seizures evoked in young APP/PS1 mice elicit premature mortality^12; 14^, likely as a result of disrupted serotonergic signaling^14^. We herein demonstrate that induction of corneal kindled seizures in aged APP/PS1 male (Figure 2G) and female (Figure 2H) mice similarly evokes a stark increase in premature mortality. Aged male and female APP/PS1 mice did not exhibit substantial differences in overall survival versus APP controls during the kindling acquisition period (Figure 2C, 2D), but once animals attained kindling criterion, there was a precipitous and substantial increase in overall mortality exclusively in APP/PS1 mice (Figure 2G; χ^2^ = 6.96, p=0.008; Figure 2H; χ^2^ = 55.0 p<0.0001). Consistent with our findings in young mice^14^, aged male and female PSEN2-N141I mice did not experience any premature mortality. Further, there was no difference in kindling or post-kindling survival between control mice of either sex, indicating the mouse strain was not responsible for increased mortality in aged APP/PS1 mice.

Established kindled seizure duration was assessed 5-7 days after all mice attained kindling criterion (Figure 2I-L). Seizure duration was modestly longer in female, but not male, PSEN2-N141I mice compared to controls (Figure 2J; p=0.016). Aged male APP/PS1 mice had longer seizure duration than APP controls (Figure 2K; p=0.0002), while female APP/PS1 mice trended towards a difference versus controls (Figure 2L; p=0.093). This suggests modest sex-related differences but an overall increase in evoked seizure severity of fully kindled APP/PS1 mice.

### Mechanistically diverse ASMs effectively reduce mean seizure severity in aged AD mice

Corneal kindling of WT mice is a moderate-throughput ASM screening model^28; 29^. We thus quantified the degree to which several mechanistically distinct ASMs could attenuate evoked seizure presentation in aged AD-associated mice (Figure 3). Age- and sex-matched AD mice exhibit notable differences in ASM activity and tolerability versus matched controls; findings that suggest that AD-related genotypes may differentially influence ASM activity beyond any variance attributable to mouse background strain alone.

**Figure 3.**
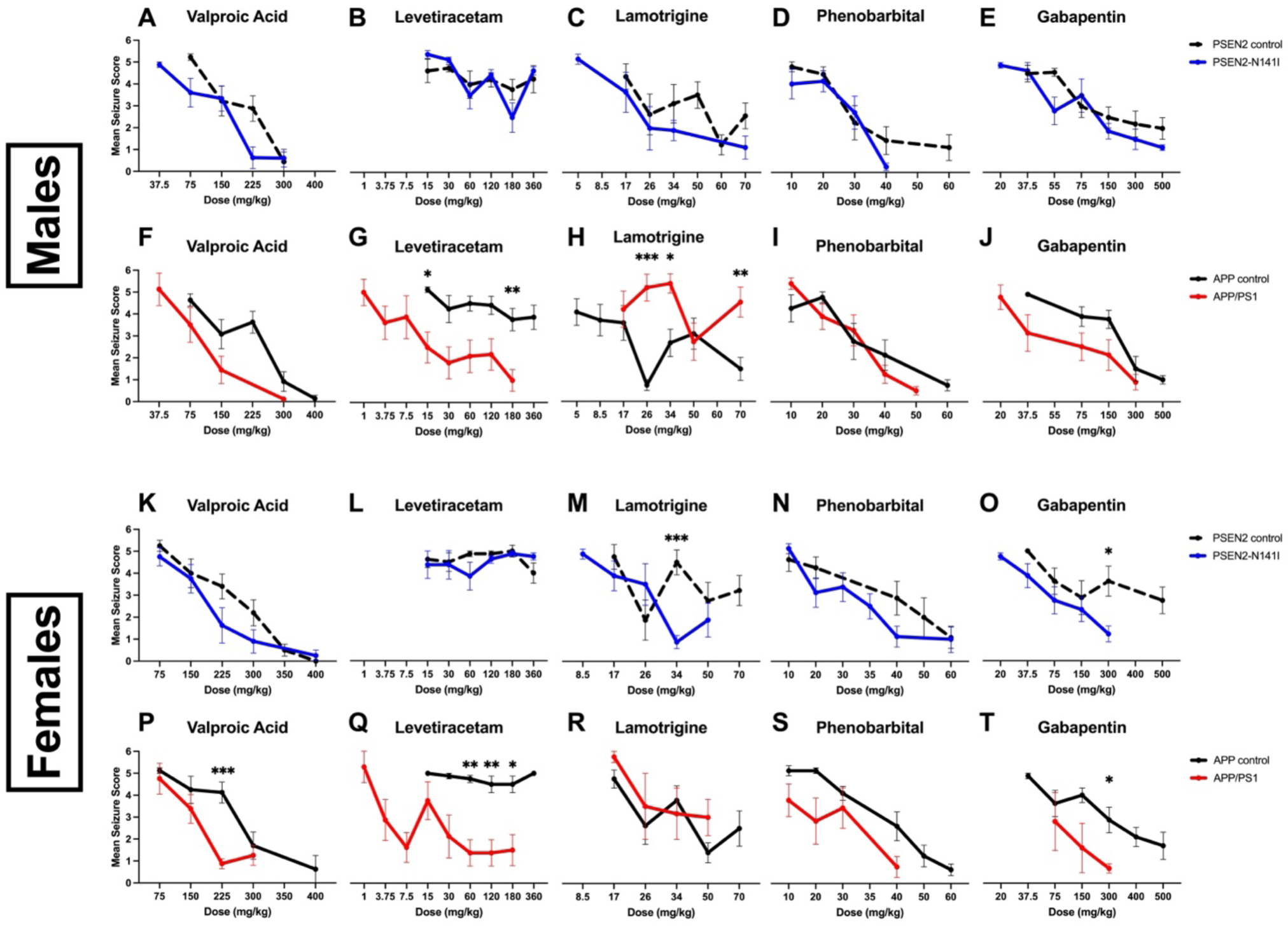
Fully kindled male and female PSEN2-N141I (blue lines) and APP/PS1 mice (red lines) were subjected to acute ASM efficacy testing over the course of 1-2 months after attaining the fully kindled state. The effects of ASM administration on mean seizure score was compared against age- and sex-matched PSEN2 control (checkered lines) and APP control littermates (black lines). Drugs tested in each genotype and sex were valproic acid, levetiracetam, lamotrigine, phenobarbital, and gabapentin all via the i.p. route. The PSEN2-N141I genotype did not significantly alter ASM efficacy in male mice as all ASMs significantly decreased seizure score in a dose related manner (A-E). Interestingly the APP/PS1 genotype in male mice significantly impacted the efficacy of multiple ASMs (F-J). LEV (G) was significantly more potent in APP/PS1 mice; doses as low as 15 mg/kg significantly differed from control mice. However, LTG (H) was significantly less potent in APP/PS1 mice, often exacerbating seizures and leading to a hyperexcitable state. There were dose-related effects in the VPA (F), PB (I), and GBP (J) testing but no significant effects of genotype. Both LTG (M) and GBP (O) had increased potency in the PSEN2-N141I female mice while VPA (K) and PB (N) showed dose-dependent efficacy. Like the male APP/PS1 data, LEV (Q) was particularly potent in female APP/PS1 compared to control mice. GBP (T) also proved to be more potent in APP/PS1 females. There were dose-dependent effects of VPA (P), LTG (R), and PB (S) in the female APP/PS1 and Tg-mice. Drugs found to demonstrate trends for dose-related seizure reduction were then subjected to dose-response quantification and ED50 calculations (Table 1).

Valproic acid (VPA) reduced seizure severity in all mice in a dose-related manner (Figure 3). APP/PS1 males showed an overall genotype effect (F_1,20_ = 6.49, p=0.019) compared to the matched APP controls, however there were no post-hoc differences between doses tested. The VPA ED50 in male APP/PS1 versus APP controls also significantly differed (Table 1). VPA was significantly more potent in PSEN2-N141I females versus controls (Figure 3K; F_1,62_ = 5.41, p=0.023), however post hoc tests again revealed no individual dose differences. Conversely, there was a dose x genotype interaction on mean seizure score in APP/PS1 and APP control females (Figure 3P; F_3,42_ = 3.64, p=0.020), with significant post-hoc seizure score differences at the 225 mg/kg dose (p=0.0005). The PI shift for APP/PS1 mice (Table 1) further confirms the increased VPA potency in APP/PS1 mice.

Levetiracetam (LEV) administration was associated with substantial genotype-related activity differences (Figure 3). While LEV administration to male PSEN2-N141I and PSEN2 controls reduced seizure score in a dose-related manner (Figure 3B; F_3,43_ = 5.96, p=0.002),genotype more significantly influenced activity of this ASM in APP/PS1 versus controls (Figure 3G; F_1,21_ = 34.15, p<0.0001). There were significant post-hoc differences at both 15 mg/kg (p=0.023) and 180 mg/kg (p=0.003). Conversely, LEV potency differed in a genotype-related manner in female APP/PS1 versus controls (Figure 3Q; F_1,14_ = 35.1, p<0.0001), with post-hoc differences at 60 mg/kg (p=0.0029), 120 mg/kg (p=0.0042), and 180 mg/kg (p=0.017). The extreme potency shift for LEV in APP/PS1 mice is further highlighted by ED50 shifts, with all other mouse strains generally showing a similar dose-response that only was notably different in APP/PS1 mice (Table 1).

Lamotrigine (LTG) administration was also associated with significant potency differences across the AD-associated genetic models (Figure 3; Table 1). There was a significant effect of genotype in male APP/PS1 mice versus APP controls (Figure 3H; F_1,67_ = 36.54, p<0.0001) with LTG proving less potent in APP/PS1 mice. There was also a significant genotype x dose interaction (F_3,67_ = 2.76, p=0.049) in male APP/PS1 and controls with post-hoc analysis indicating that the 26 mg/kg (p=0.0003), 34 mg/kg (p=0.011), and 70 mg/kg (p=0.008) doses were less potent in APP/PS1 mice. The LTG ED50 was 28.1 mg/kg [11.6-51.7] for APP control males, but not even high dose (70 mg/kg) LTG that can adversely affect motor coordination in WT mice^15; 24^, could suppress seizures in aged male APP/PS1 mice (Table 1). LTG administration to female PSEN2-N141I mice conferred a significant dose x genotype interaction (Figure 3M; F_3,42_ = 5.47, p=0.003), with 34 mg/kg (p=0.0007) differentially reducing mean seizure score in PSEN2-N141I versus controls.

Phenobarbital (PB) reduced seizure severity in a dose-related manner in all genotypes, regardless of sex (Figure 3). Further, there was a significant effect of APP/PS1 genotype in female mice (Figure 3S; F_1,39_ = 11.41, p=0.002) with PB proving more potent than in controls. The PB ED50 and PI were also determined for all male and female kindled mice and with no significant differences in PB potency or tolerability across any genotypes or sexes (Table 1), consistent with the broad anticonvulsant activity of this compound in rodent epilepsy models.

Gabapentin (GBP) is a commonly prescribed ASM in older adults with epilepsy^30; 31^. Male PSEN2-N141I and APP/PS1 mice and matched controls were sensitive to GBP in a dose-related manner, however seizure control only significantly differed between APP/PS1 and matched controls (Figure 3J; F_1,16_ = 9.76, p=0.007). However, the GBP ED50 did not differ across any male mice (Table 1). There was a significant genotype effect between female PSEN2-N141I and matched controls (F_1,44_ = 6.75, p=0.013; Figure 3O), with post hoc differences at 300 mg/kg GBP (p=0.03). A similar genotype effect occurred in female APP/PS1 and controls (F_1,12_ = 6.48, p=0.026; Figure 3T), with post hoc differences at 300 mg/kg (p=0.019). These ED50s revealed significantly greater GBP potency in female AD-associated mice versus their controls (Table 1).

### AD genotype influences change in acute seizure suppression by ASMs

We also assessed seizure score changes between the testing session 24-hours before each ASM administration (Figure 4). Only mice that had a Racine stage 4+ seizure received an ASM the following day. ASM doses that more significantly reduced seizure severity in the AD mouse are presented on the right side of the graph; ASM doses that more significantly reduced seizure severity in control lines are on the left. Response to LTG and GBP administration most meaningfully differed between PSEN2-N141I and controls (Figure 4), whereas response to VPA and LEV administration was most notably shifted in APP/PS1 versus controls (Figure 4). These data further support the hypothesis that AD-related genotypes differentially influence ASM activity, underscoring a possible opportunity to precisely tailor ASM use in clinical AD.

**Figure 4.**
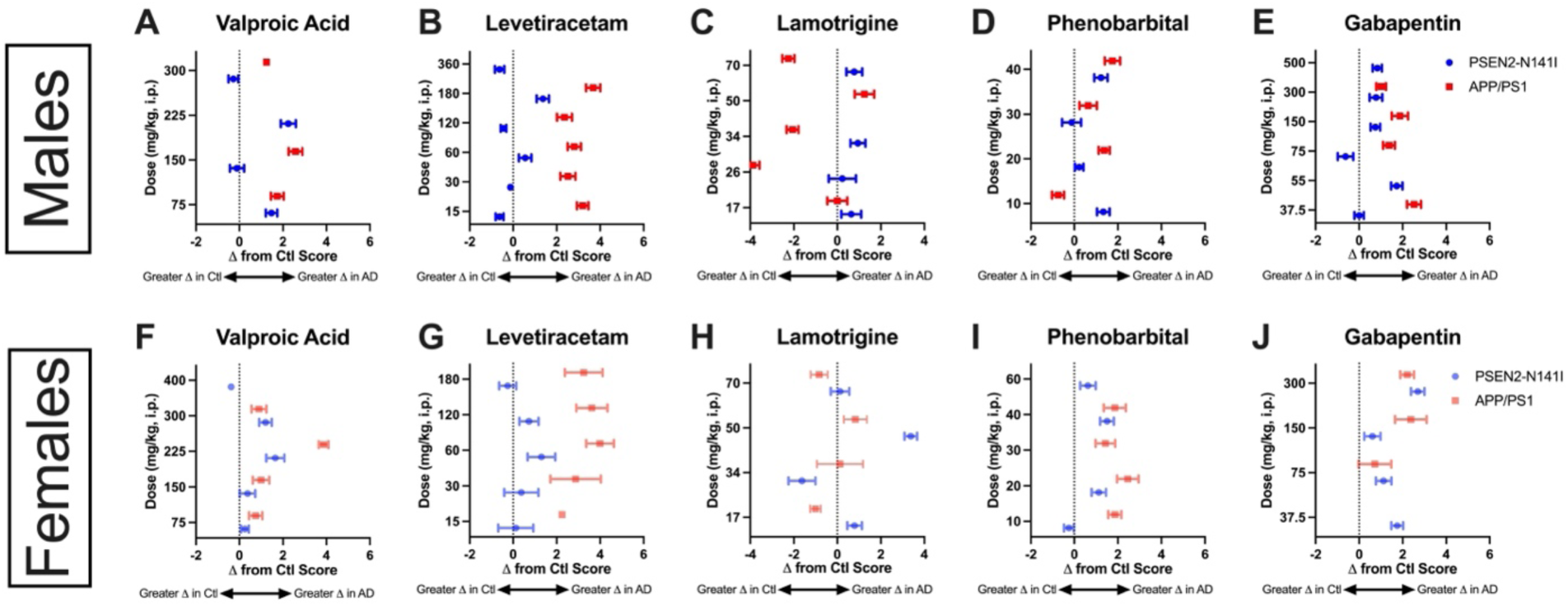
The shift in baseline seizure score versus the ASM testing session 24 hours later was compared across all animals and genotypes to determine the magnitude of ASM efficacy. Drugs that more meaningfully reduced seizure score in control animals are on the left-hand side of the plot whereas drugs that more meaningfully changed seizure score in the AD genotypes are presented on the right-hand side of the plots. Importantly, most ASMs more substantially reduced mean seizure score from baseline scores in AD-related genotypes, regardless of sex.

### Aβ accumulation is only evident in aged APP/PS1 mice

Finally, we investigated whether corneal kindling in aged AD or WT mice accelerated the accumulation of Aβ plaques. No study has yet demonstrated whether aged PSEN2-N141I mice exhibit any pathological Aβ accumulation, despite evidence that other PSEN2 transgenic mouse models do not show Aβ accumulation unless crossed to an APP-overexpressing line^32; 33^. Aged PSEN2-N141I mice, regardless of kindling status, did not demonstrate Aβ plaques at 8-15 months-old (Figure 5A-D).

**Figure 5.**
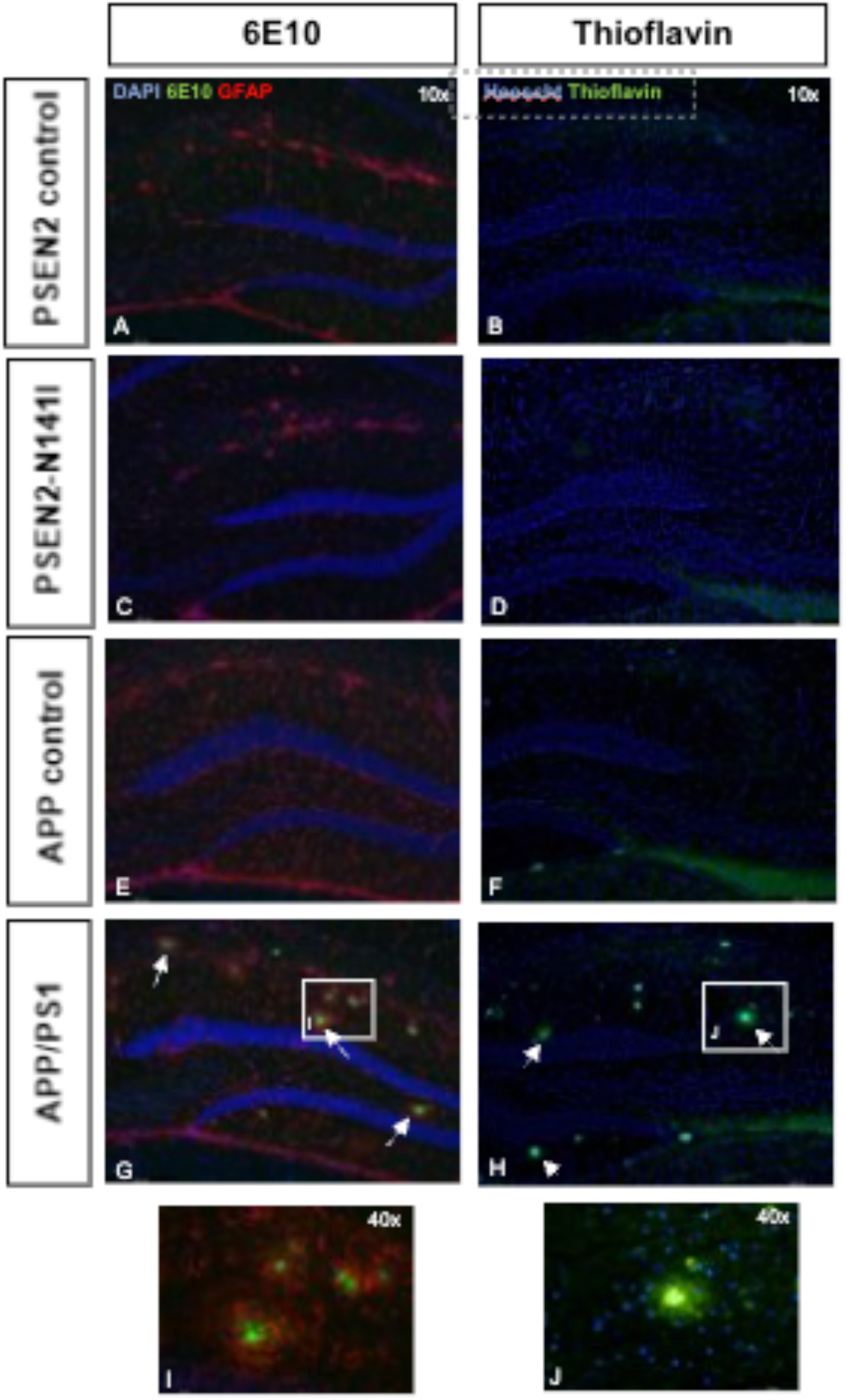
The presence of Aβ aggregates was assessed in both sexes in PSEN2 control, PSEN2-N141I, APP control, and APP/PS1 mice. Qualitative assessment of Aβ accumulation as assessed by both 6E10 immunofluorescence (A, C, E, and G) and Thioflavin S staining (B, D, F, and H) in aged, fully kindled APP/PS1 and PSEN2-N141I mice that were not subjected to anticonvulsant efficacy testing with candidate ASMs. Further, tissues were labeled for the astrocytic marker, GFAP, to demonstrate widespread neuroinflammation in kindled animals. Only aged APP/PS1 mice (G, H) demonstrate widespread accumulation of Aβ plaques, which is absent in APP control mice, as well as kindled PSEN2-N141I and PSEN2 control mice. Notably, astrocytes (I) were evident surrounding Aβ plaques in APP/PS1 mice. J) Representative higher magnification of Thioflavin S labeling (40x).

We have shown that fully kindled young adult (2-3 months old) APP/PS1 and 3xTg mice do not have Aβ plaques^14; 18^. We now demonstrate that corneal kindling aged APP/PS1 mice does not qualitatively increase Aβ plaque deposition (Figure 5). Only APP/PS1 mice had Aβ plaques (Table 2). Further, Aβ aggregates were encircled by GFAP-positive labeling (Figure 5I), indicative of reactive astrogliosis.

## Discussion

Our present study extends earlier work to clearly demonstrate that investigator-induced neuronal hyperexcitability in aged mice with AD-related genotypes is not universally detrimental and that genotype-related interactions additively worsen disease burden. We demonstrate several novel findings. First, AD-associated risk factors differentially influence susceptibility to 60 Hz corneal kindled seizures in late life, representative of clinical AD. These data comprehensively extend our earlier work with young, pre-symptomatic APP/PS1^14; 18^, 3xTg mice^18^, PSEN2-N141I^14^, PSEN2 KO mice^15; 19^. Notably, we herein establish that the PSEN2-N141I variant prevents the formation of a fully kindled state even at late ages in a sex-related manner, whereas aged APP/PS1 mice are not more susceptible to 60 Hz corneal kindling (Figure 1). These data cumulatively suggest that biological sex, age, and AD-associated genotype greatly influence susceptibility to form a hyperexcitable neuronal network. Second, our study reveals that AD-associated risk genes differentially influence seizure-related chronic outcomes. Aged APP/PS1 mice were at a substantially heightened risk of seizure-induced premature mortality; an effect not observed in PSEN2-N141I or control mice (Figure 2). These findings in aged APP/PS1 mice match our earlier findings in young APP/PS1 mice^14; 18^ and further reveal that evoked neuronal hyperexcitability in an amyloidogenic background can worsen disease course. Lastly, we now show that AD-associated mouse lines are sensitive to a diversity of mechanistically distinct ASMs with genotype and sex markedly influencing ASM potency and tolerability. The pharmacological potency and tolerability profiling of commonly prescribed ASMs in aged AD mouse models reveals possible opportunities to more precisely manage seizures in adults with AD. Altogether, this study confirms and extends evidence of the adverse functional consequences of chronic focal seizures in AD models, particularly in models with high Aβ levels.

One important finding is the clear dissociation between Aβ-dependent and Aβ-independent influences on seizure-induced premature mortality. Prior studies of spontaneous seizures in aged AD models suggested that epileptiform activity secondary to Aβ accumulation was responsible for the precipitous mortality in APP/PS1 mice^34^. Although young, presymptomatic APP/PS1 mice similarly succumb to seizure-induced premature mortality^14^, the present study in appropriately aged mice with two different AD-related genotypes now clearly confirms that evoked convulsive seizures only elicit premature mortality with Aβ processing deficits (Figure 2). Specifically, only APP/PS1 mice were impacted, being most consequential in aged female APP/PS1 mice. People with AD and comorbid seizures experience hastened mortality versus individuals without seizures^35; 36^. Indeed, serotonergic system dysfunction, implicated in sudden unexpected death in epilepsy (SUDEP)^37^, occurs in young, kindled APP/PS1 mice^14^ and serotonin synthesis deficits increase mortality in APP/PS1 mice^34^. Our study thus emphasizes the urgent need to define how seizures negatively influence survival, and the relationship between Aβ processing and serotonin signaling.

Evoked seizures in AD-related mice can assess the pathological and functional impacts of chronic seizures and rapidly screen for potential novel treatments^15^. Corneal kindling generates many mice with uniform seizure history for sufficiently powered pharmacological studies to rigorously quantify ASM potency and tolerability^28; 38^. Unlike the long-duration recording of spontaneous electrographic seizures in single-housed mice^39–42^, corneal kindling employs socially-housed mice^43; 44^. Applying this seizure induction paradigm to mice with discrete AD-associated genotypes facilitates targeted assessment of biological heterogeneity of seizure-induced functional decline^16^. It also establishes that the tested ASMs generally control seizures in mice with AD-associated backgrounds in a dose-related manner, therefore any cognitive benefits uncovered in prior studies may not be solely due to specific ASMs^39; 45–49^, but rather to the general suppression of neuronal hyperexcitability^2; 21; 50^.

We uncovered unique pharmacological potency and tolerability differences with discrete ASMs in AD-related models, suggesting a precision medicine-based strategy for seizures in AD. For example, the benefit of LEV in patients with AD^21; 50^ and MCI^51^ is being actively investigated. However, our present data and studies by others do not support that LEV is the only ASM that can improve seizure burden. Older adults with epilepsy are commonly prescribed LTG, and this drug has a high retention rate in this population^30^. Indeed, Na+ blockers can improve cognitive function and neuropathological hallmarks of AD^52^. Aβ_1-42_ itself may modulate voltage-gated sodium channel Nav1.6 expression, thereby promoting neuronal hyperexcitability^53^. Aβ plaques were absent in PSEN2-N141I mice (Figure 4); it is thus possible that PSEN2-N141I mice were more sensitive to LTG because of conserved Nav1.6 expression due to the absence of Aβ accumulation. Further, GBP was also effective and well-tolerated across AD-genotypes, findings that have yet to be reported elsewhere in any AD-related model. ASM selection should be nonetheless carefully considered in clinical practice; despite our present evidence that VPA and PB exerted consistent dose-related seizure control, both ASMs can worsen cognitive function in people with AD^54^ Moreover, PB was associated with beneficial dose-related effects on seizure severity, yet Cumbo and Ligori reported that PB evokes negative cognitive effects in AD.^55^. Nonetheless, our findings illustrate the divergent biological contributions of presenilins versus Aβ on AD-related seizure susceptibility and highlight that evoked seizures in AD-associated rodents can be feasibly uncover disease-modifying treatments.

Kindling rodents with AD-related genotypes is increasingly applied to understand how seizures in AD may influence disease processes and cognitive function^14; 19^. In contrast to 6 Hz corneal kindling in two AD-associated models^12^, our present data illustrate that chronic kindled seizures in AD-associated models are not drug-resistant, but that ASM resistance may instead depend on the specific seizure induction protocol (e.g., 60 vs 6 Hz)^12^. The 6 Hz corneal kindling paradigm itself evokes a highly drug-resistant seizure^56^, whereas 60 Hz corneal kindling is generally not treatment-resistant without additional manipulation^29; 57^. Our approach therefore allows for discrete pharmacological profiling in an otherwise drug-sensitive chronic focal seizure model in aged rodents to more comprehensively assess how AD-related genotypes influence ASM activity. Such information may inform precisely tailored seizure management in clinical AD. These present data conclusively demonstrate that seizure control is attainable in AD-associated models and that AD-related factors (i.e. Aβ) critically and differentially influence ASM activity. Ultimately, this study illustrates that the 60 Hz corneal kindling paradigm with aged AD-associated mice may uncover novel drugs for seizures in clinical AD.

## Acknowledgements/Conflicts/Funding Sources

The authors are grateful for technical assistance from Rami Koutoubi. None of the authors have any conflicts to disclose. This work was supported by an American Epilepsy Society Junior Investigator Award and NIA R01AG067788 to MBH.

## Abbreviations

ASM: antiseizure medicine
AD: Alzheimer’s disease
WT: wild type
APP: amyloid precursor protein
PSEN1: presenilin 1
PSEN2: presenilin 2
Aβ: amyloid beta
KO: knock out
PBS: phosphate buffered saline

## Supplemental Tables and Figures

**Supplemental Figure 1.**
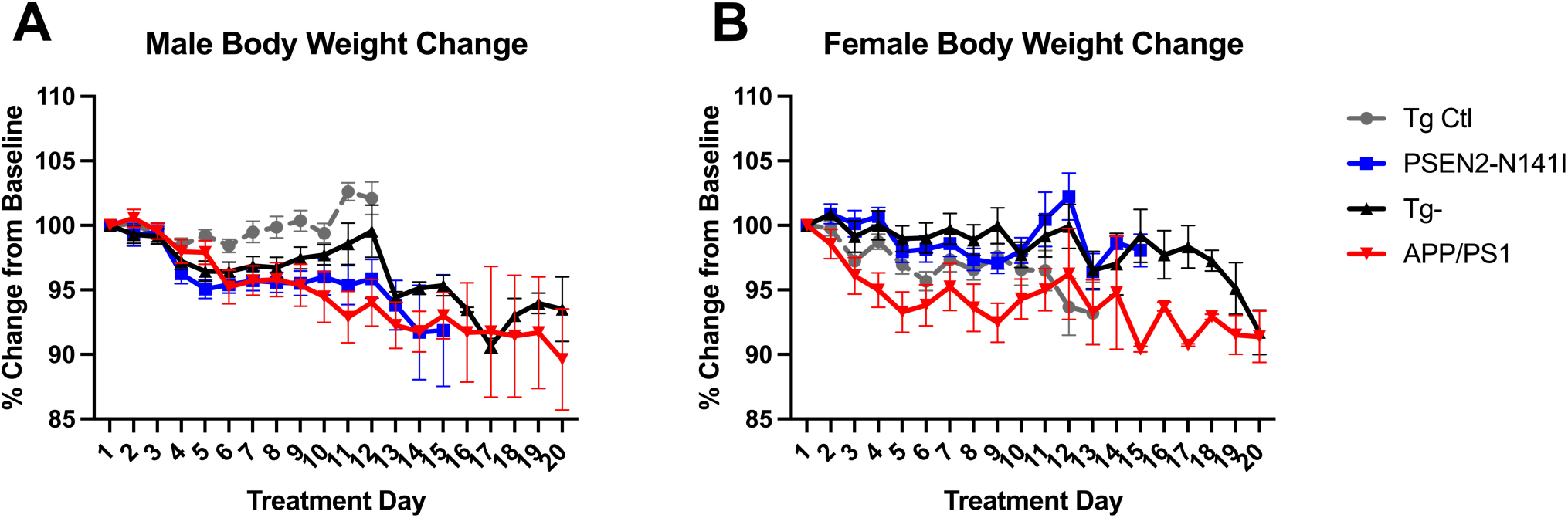
Body weight change during the repeated ASM testing period after mice attained kindling criterion. Body weights were obtained on each drug administration day, with testing occurring no more than twice per week. There were no significant differences in body weight change from baseline over the course of the ASM testing period.

**Supplemental Table 1.**
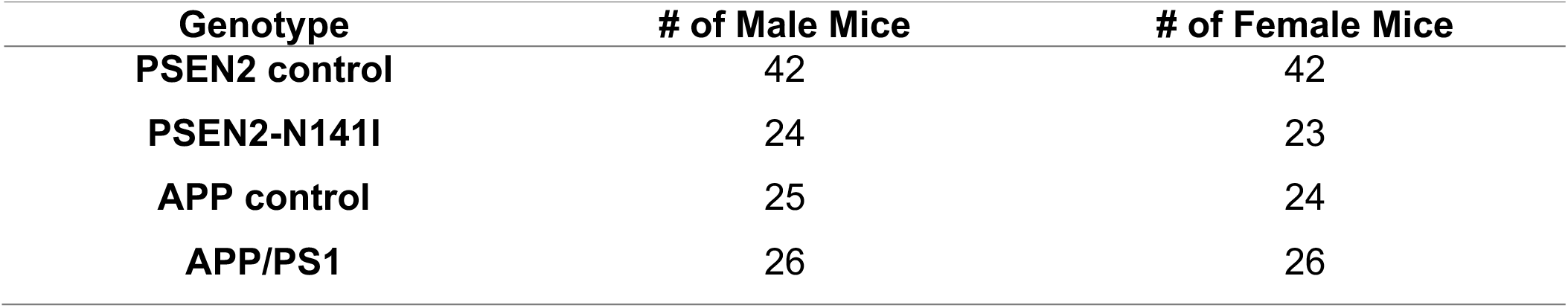
Total number of both male and female mice from each genotype during the pharmacology testing.

