## Supplemental Table and Figure for "Alzheimer’s disease-associated genotypes differentially influence chronic evoked seizure outcomes and antiseizure medicine activity in aged mice"

**Supplemental Tables and Figures.**

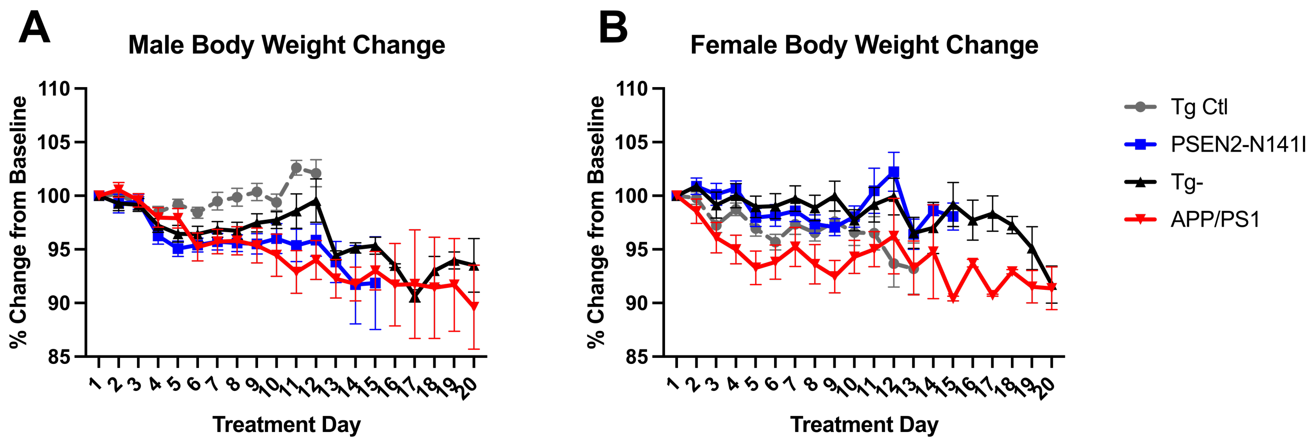


**Supplemental Figure 1.** Body weight change during the repeated ASM testing period after mice attained kindling criterion. Body weights were obtained on each drug administration day, with testing occurring no more than twice per week. There were no significant differences in body weight change from baseline over the course of the ASM testing period.

**Supplemental Table 1**. Total number of both male and female mice from each genotype during the pharmacology testing.

| Genotype | # of Male Mice | # of Female Mice |
|---|---|---|
| **PSEN2 control** | 42 | 42 |
| **PSEN2-N141I** | 24 | 23 |
| **APP control** | 25 | 24 |
| **APP/PS1** | 26 | 26 |
